# Detecting strawberry cultivar misidentification in the Philippines using single nucleotide polymorphism markers from the anthocyanin reductase gene

**DOI:** 10.1101/2020.02.02.922369

**Authors:** Nadine Adellia Ledesma, John Mark Matulac, Jesus Emmanuel Sevilleja, Maria Luisa Enriquez

## Abstract

**BACKGROUND:** Commercial strawberry production in the Philippines is done by small-holder farmers in La Trinidad, Benguet, where the climate is conducive for optimal growth of this temperate crop. However, these farmers are not cognizant of the importance of proper cultivar identification, particularly during runner propagation, distribution, and transplanting. Thus, there is a high likelihood that misidentification of commonly grown cultivars has taken place.

**OBJECTIVE:** The study aimed to develop single nucleotide polymorphism markers and use them to detect possible misidentification among strawberry cultivars.

**METHODS:** Leaf samples from several cultivars were obtained from farmers and the germplasm collection of a local university in La Trinidad, Benguet, Philippines. Expressed sequence tags from the *ANR* gene were screened for putative SNPs. Eleven SNP markers were developed and used to discriminate among the collected samples.

**RESULTS:** The SNP markers grouped the cultivars into five genotypic clusters with seven distinct genotypic identities. Clustering analysis revealed inconsistencies between the farmers’ identification and the molecular classification. ‘Sweet Charlie’ samples were assigned to four genotypic clusters and ‘Strawberry Festival’ samples were grouped into three separate clusters.

**CONCLUSION:** There is a high probability that cultivar misidentification has indeed occurred. The molecular markers developed in this study could assist in future cultivar verification efforts, germplasm management, and breeding programs.

## 1. Introduction

The cultivated strawberry (*Fragaria* × *ananassa* Duch.) is a temperate fruit crop that has seen a 230% increase in worldwide production in the past three decades. Seventy-seven countries produced a combined 9.2 million tons of the fresh fruit in 2017 [1]. While strawberry production has traditionally been associated with temperate countries, strawberries can nonetheless grow in the colder mountainous areas of the tropics [2]. In Asia, the Philippines is the only tropical country that grows strawberries on a commercial scale. The country produced 819 tons of fresh strawberries in 2017 [1].

Ninety-five percent of strawberry production in the Philippines is located in the high-altitude municipality of La Trinidad, Benguet (16°28’N 120°35’E, 1300 m). The region is conducive for strawberry growth and development because of the relatively mild temperatures throughout the year. Much of the strawberry production is done by small-holder farmers in an area called the Strawberry Farm. Strawberry cultivars that have been grown in the area include ‘Sweet Charlie’, ‘Chandler’, and ‘Whitney’ from the United States, and ‘Toyonoka’ and ‘Nyoho’ from Japan. Currently, the Strawberry Farm mainly produces ‘Sweet Charlie’. Many of the strawberry farmers use runners obtained from several generations of vegetative propagation, often swapping with or providing planting materials to neighboring farmers. This exchange of runners has led to a potentially serious problem of cultivar misidentification. In addition, different cultivars are usually grown in the same plot or in nearby plots to obtain runners for the succeeding growing season, and farmers have not always been concerned about mixing cultivars, much less properly labeling them when propagating, transferring, and planting [3]. To exacerbate matters, whenever new planting materials arrive, farmers do not follow proper naming conventions in identifying cultivars, often naming them based on the country of origin. For instance, ‘Sweet Charlie’ planting materials obtained from Argentina are named ‘Sweet Charlie’ Argentina and those from Washington, USA are named ‘Sweet Charlie’ Washington.

Cultivar misidentification has been known to occur during asexual propagation of strawberry plants in nurseries in the United States. These nurseries are the sources of planting materials for commercial growers [4,5]. Although morphological traits can be used to differentiate strawberry cultivars, this technique has been found to be inconsistent because the morphological differences are often small [6] and ambiguous [7], especially during the early vegetative stage and if two cultivars have a similar genetic lineage. Furthermore, phenotypic alterations due to prevailing environmental conditions can also lead to cultivar misidentification (e.g., plant height, fruit size, and fruit sweetness are affected by nutrient availability and temperature) [7].

Providing genetically correct planting stock is crucial in commercial strawberry production because growers select specific cultivars for desirable traits that match customer preference and the production capacities and limitations of the growers (e.g., yield, fruit size, and pest and disease resistance) [8]. Thus, cultivar verification should be an essential step for farmers. It is also necessary for certification programs and for the protection of the rights of plant breeders [9]. Cultivar identification at the molecular level could solve this problem not just for the strawberry but for other economically important crops that may be difficult to distinguish morphologically or for those whose genetic lineage is unknown.

There have been several attempts to differentiate strawberry cultivars at the DNA level using various molecular markers. Reported methods include randomly amplified polymorphic DNA (RAPD) markers [7,10–12], amplified fragment length polymorphism (AFLP) markers [13–15], cleavage amplified polymorphisms (CAPS) [9], simple sequence repeats (SSR) [16,17], and combinations of these markers [18–20]. A more recent molecular method of cultivar discrimination is the use of single nucleotide polymorphism (SNP) markers.

The methods for SNPs discovery and validation have been perfected for several diploid crop species, but the same cannot be said for many of the world’s most important crops (e.g. wheat, potato, sugarcane, and cotton) because they are polyploids. SNPs discovery in these crops is mainly hindered by the complexity of their genome [21–23]. The larger number of homologues has also posed challenges in the correct identification of SNPs [23].

Such are the challenges being faced by the cultivated strawberry, an allo-octoploid (2*n* = 8*x* = 56) whose subgenome composition has not been fully elucidated [22]. This is evident from the limited number of studies published on SNP markers for this crop, although successful SNP discovery has been made in the diploid *Fragaria vesca* [24,25].

Several strategies for SNPs discovery and validation have been reported in other polyploid crops [21]. One such method is the use of high-resolution melting curve analysis (MCA) using fluorescent dyes following real-time PCR, which was employed in the autopolyploid alfalfa [26–28]. The advantage of MCA is that it provides a single-step, closed-tube method that do not require time-consuming confirmatory steps such as gel electrophoresis [29]. The tandem use of real-time PCR and MCA can thus simplify SNP detection, quantification, and genotyping [30]

In the Philippines, there have been no attempts to verify strawberry cultivars, either at the morphological or molecular level. If farmers have not been careful during their asexual propagation of runners from cultivars growing near each other, then there is a high probability that a mix-up may have occurred, either during the runner harvest stage or the transport/distribution stage. Therefore, this study aimed to explore the possibility of cultivar misidentification in the main strawberry producing area in the Philippines by employing molecular techniques. The objectives were (1) to develop a relatively inexpensive, simple, fast, sensitive, and high-throughput, single-tube assay for SNP detection in strawberry cultivars being grown in the Philippines; and (2) use the developed SNP markers to discriminate amongst the cultivars and determine if any misidentification has indeed occurred.

## 2. Materials and methods

### 2.1. Mining of putative SNPs from nucleotide sequences and ESTs

All available nucleotide sequences of *Fragaria* that code for a trait that affects fruit quality were downloaded and screened from the NCBI Genbank (http://www.ncbi.nlm.nih.gov, March 20, 2016). A BLAST search of the target sequences was performed and all available ESTs that matched the sequence were downloaded in FASTA format. Assembly of ESTs were then limited to the *ANR* gene. Assembly of the ESTs were done using the CAP3 sequence assembly program (http://doua.prabi.fr/software/cap3). A contiguous sequence/contig (consensus sequence) was generated and aligned with and a reference genomic sequence using MEGA 6.0 to identify exonic and intronic regions and candidate target regions for putative SNP discovery (sequence <200bp and located between relatively conserved end regions).

### 2.2. Primer design

Primers were designed using Primer3 (http://bioinfo.ut.ee/primer3/) to generate a list of candidate primer pairs. Viability of the primer pairs for successful amplification was assessed using NetPrimer Primer Analysis (http://www.premierbiosoft.com/netprimer/). Criteria for selection was based on difference in melting temperature <5°C, GC content ~50% and probability of cross dimer, self-dimer and hairpin structure formation ΔG>-9.0 kcal. A BLAST search of the target sequence was also done for verification of specificity to the genus *Fragaria*.

A total of five primer pairs targeting the same region of the *ANR* gene was designed (Table 1). Primer Pairs A, B and C had Tm difference of ≈ 5°C and GC% = 50±15%. Self-dimers, hairpin structures and cross-dimers were present, but all were within the acceptable values of ΔG ≥ −9.0. Primer Pair D and E had Tm difference of ≈ 5°C, GC% = 50±3%, and self-dimer and hairpin ΔG = 0. Although there was an instance of cross dimers at ΔG = −4.9 for Primer D and ΔG = −6.03 for Primer E.

**Table 1.**
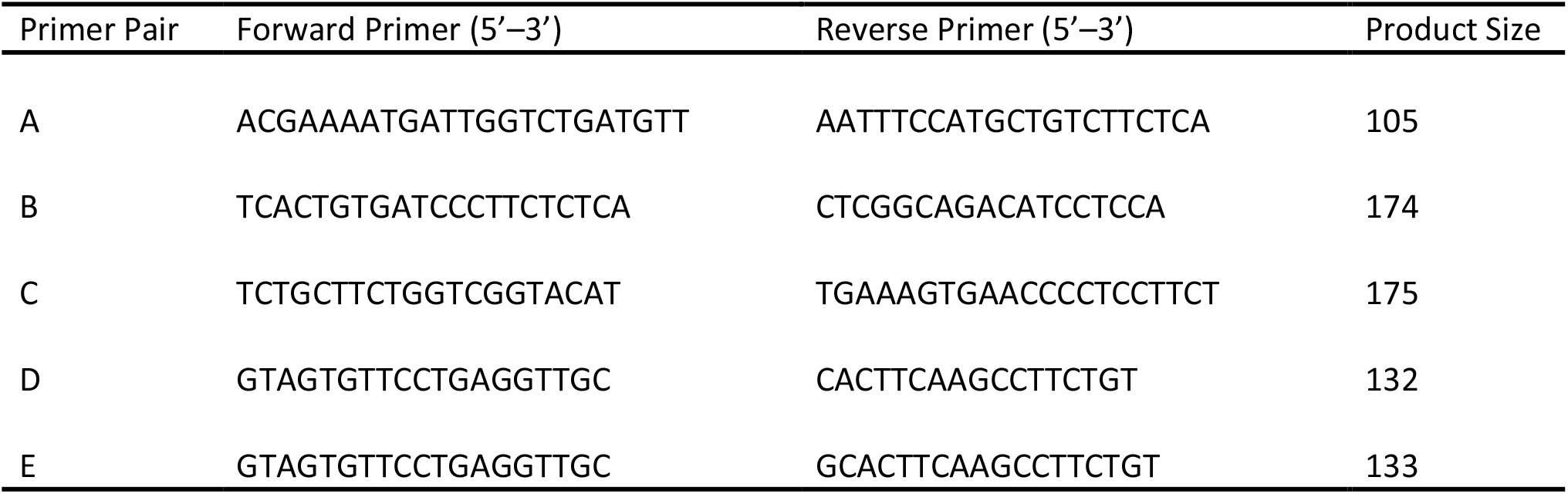
Designed primer pairs for the target region in the *ANR* gene

### 2.3. Field collection of strawberry cultivars

Plant specimen collection was conducted from April 2016 to February 2017 in La Trinidad, Benguet, Philippines. Collection of leaf samples from commercially grown cultivars were limited to the Strawberry Farm and surrounding areas. Fully expanded trifoliate leaves were cut off and placed in plastic air-tight bags. All leaf samples were stored at −20°C until DNA extraction. A total of twenty (20) specimens were collected (Table 2). The names of the cultivars were provided by the farmers who were growing them, which they identified based on the cultivar and supposed country from which the original planting materials were obtained.

**Table 2.** List of strawberry cultivars collected in La Trinidad, Benguet with GPS coordinates.

| Sample | Label | Coordinates |
| --- | --- | --- |
| 1. | 'Harunoka' | 16.455278, 120.584955 |
| 2. | 'Summer Princess' 2 | 16.457357, 120.583056 |
| 3. | 'Summer Princess' 3 |  |
| 4. | 'Winter Dawn' 2 | 16.457439, 120.582155 |
| 5. | 'Winter Dawn' 3 |  |
| 6. | 'Sweet Charlie' 1 | 16.455083, 120.582670 |
| 7. | 'Sweet Charlie' 2 | 16.456040, 120.582391 |
| 8. | 'Sweet Charlie' 3 | 16.456246, 120.583239 |
| 9. | 'Sweet Charlie' 4 | 16.457532, 120.585535 |
| 10. | 'Sweet Charlie' 5 | 16.457439, 120.582155 |
| 11. | 'Sweet Charlie' 6 | 16.452253, 120.576683 |
| 12. | 'Sweet Charlie' California | 16.453467, 120.583013 |
| 13. | 'Sweet Charlie' Washington 1 | 16.450051, 120.591950 |
| 14. | 'Sweet Charlie' Washington 2 | 16.453467, 120.583013 |
| 15. | 'Sweet Charlie' Argentina 1 | 16.450051, 120.591950 |
| 16. | 'Sweet Charlie' Argentina 2 |  |
| 17. | 'Sweet Charlie' Argentina 3 | 16.453467, 120.583013 |
| 18. | 'Strawberry Festival' 1 | 16.450051, 120.591950 |
| 19. | 'Strawberry Festival' 2 |  |
| 20. | 'Strawberry Festival' 3 | 16.453467, 120.583013 |

Note: \* Cultivar names were provided by the farmers. Numbers after each cultivar were added by the authors to indicate leaf samples collected from different plants.

**Table 2.** Specificity of Primers C, D, and E in amplifying specific strawberry samples from the Strawberry Farm, La Trinidad, Benguet, Philippines

| Sample No. | Primer |  |  |
| --- | --- | --- | --- |
|  | C | D | E |
| 4 | ✓ |  | ✓ |
| 5 | ✓ | ✓ |  |
| 7 | ✓ | ✓ | ✓ |
| 9 |  | ✓ | ✓ |
| 11 |  | ✓ | ✓ |
| 12 | ✓ |  | ✓ |
| 13 |  | ✓ | ✓ |
| 14 |  | ✓ | ✓ |
| 16 |  | ✓ | ✓ |
| 17 |  | ✓ | ✓ |
| 18 | ✓ | ✓ | ✓ |
| 19 | ✓ |  |  |
| 20 | ✓ | ✓ | ✓ |

### 2.4. DNA extraction and Real-Time PCR

Young leaves from each of the collected specimens were lyophilized in liquid nitrogen then ground into a fine powder with a mortar and pestle. DNA was extracted from the powdered samples using the NucleoSpin™ Plant II (Macherey-Nagel, Germany) following the protocol specified in the kit, and stored at −20°C prior to PCR analysis.

Extracted DNA samples were amplified by real-time PCR using the designed primers. The 30μL initial PCR reaction was a mixture of the following: 15μL SSoAdvanced™ Universal SYBR Green Supermix, 6μL DNA, 6μL nuclease-free water, 1.5μL Forward primer and 1.5μL Reverse primer. Using the Bio-Rad^®^ CFX96 Real-time PCR system (Bio-Rad®, USA), cycling parameters were as follows: Polymerase activation and DNA denaturation at 98°C for 5 min and 40 cycles at 98°C for 15 sec denaturation, optimum temperature for specific primer pair (°C) for 60 sec annealing/extension. A replicate run of all samples was also done and subjected to melting curve analysis (65–95°C, 0.5°C at 2–5 sec/step) to verify if the amplicon products were pure and single.

Post-amplification melting-curve analysis is a simple, straightforward way to check real-time PCR reactions for primer-dimer artifacts and to ensure reaction specificity. This method revealed that Primer Pairs A and B were not suitable for target sequence amplification due to their lack of reaction specificity. Thus, only Primer Pairs C, D, and E were used in the study. The PCR amplicons were further subjected to gel electrophoresis to verify the presence of a single band as an added visual confirmatory step.

For DNA sequencing, PCR amplicons were sent to Macrogen, Inc. (South Korea) for processing as follows: the amplicons were first cleaned using the ExoSAP-IT PCR Clean-up Kit following the manufacturer’s instructions (USB Corp., Cleveland, OH, USA). The purified amplicons were Sanger-sequenced using the BigDye Terminator v3.1 Cycle Sequencing Kit on both forward and reverse strands then analyzed using the ABI PRISM 3730XL Analyzer (Applied Biosystems, Foster City, CA, USA).

The sequence data were aligned in MEGA 6.0 to screen for any differences in sequence information such as base changes or insertions/deletions and to resolve possible errors in sequencing and flanking ends. The sequences were further analyzed in BLAST to verify specificity of the sequences to the *ANR* gene. The total number of SNPs and SNPs distribution (i.e., transversions, transitions, and insertions/deletions) were also determined.

### 2.5. Cluster analysis

The SNP markers were then checked for their suitability in cultivar discrimination. Phylogenetic analysis of SNP data was done using the Neighbor–Joining method on MEGA 6.0 to cluster closely related cultivars based on SNP characteristics.

## 3. Results and discussion

### 3.1. Data mining and assembly of strawberry ESTs

A BLAST search in NCBI (http://www.ncbi.nlm.nih.gov) for the genus *Fragaria* revealed 60,035 ESTs, 10,988 ESTs for *Fragaria* × *ananassa*, and eight ESTs for *Fragaria* × *ananassa* ‘Sweet Charlie.’ ‘Sweet Charlie’ was specifically searched because it is the most widely grown cultivar at the Strawberry Farm. Among the eight candidate ESTs found for ‘Sweet Charlie,’ the anthocyanidin reductase gene (*ANR*, AN: JX271492.1) was selected for putative SNP discovery. The BLAST search for *ANR* resulted in 24 EST sequences. Assembly of these ESTs produced a 1261-bp long contig. This contig was then aligned with the reference genomic sequence (DQ664193.1 *Fragaria* × *ananassa* anthocyanidin reductase (*ANR*) gene complete). The alignment revealed two introns and three exons within the *ANR* sequence (Fig. 1). A sequence of about 200 bp with relatively conserved flanking regions in Exon 3 was targeted as the template to design the primers.

**Figure 1.**
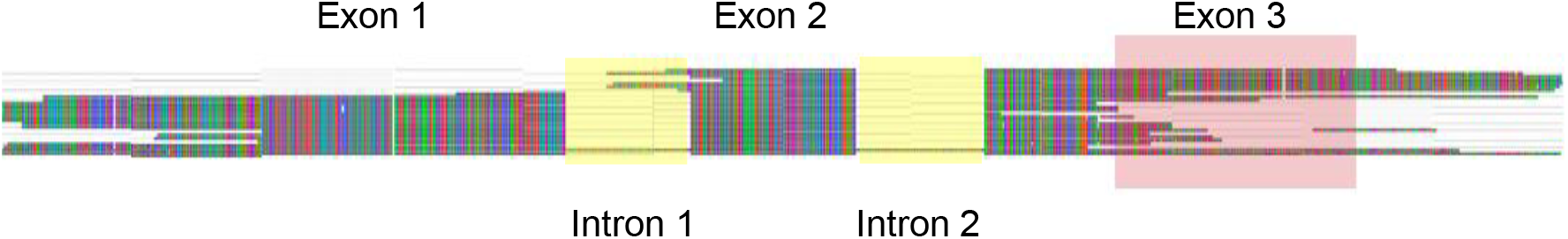
Assembly of Strawberry *ANR* ESTs. The highlighted red region represents the target sequence, yellow region represents the introns.

The *ANR* is a gene of interest because there is a high probability that this region of the genome might have undergone genetic modification from breeding programs due to its involvement in proanthocyanidin (PA) biosynthesis. This gene regulates the conversion of PAs, which are known flavonoids that affect the yield, survivability, forage traits and healthiness of grown crops [31]. The gene is highly expressed during fruit development where PAs function for protection against biotic and abiotic stress and have great economic significance due to their influence on the nutritional content of crops as a rich source of antioxidants [31,32]. These properties make the *ANR* a useful gene for marker-assisted selection (MAS) and cultivar identification. Whitaker highlighted the need to develop molecular markers for fruit quality traits in the cultivated strawberry [4], a task which has not been the major focus of breeding programs as of late. The SNP markers for the *ANR* gene developed in this study could thus be a starting point for MAS or pedigree-based analysis in this important crop.

### 3.2. Real-time PCR amplification and sequencing

Only 14 out of the 20 strawberry samples collected were successfully amplified by the three primers. Real-time PCR amplification and subsequent melting curve analysis revealed clean, unambiguous curves for the samples (Fig. 2). Agarose gel electrophoresis further confirmed the successful amplifications (data not shown).

**Figure 2.**
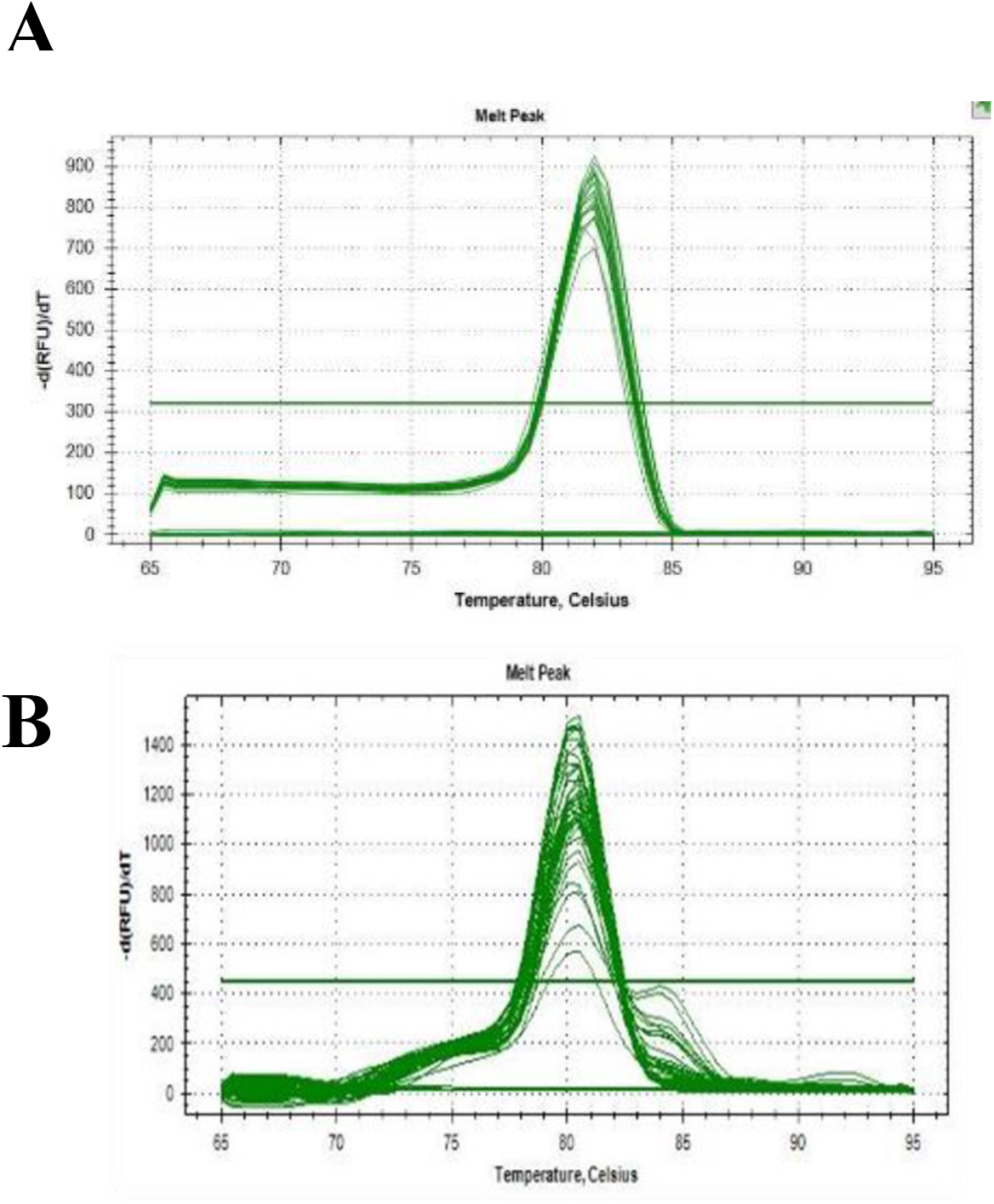
Melting curve analysis using different primer pairs to check specificity of designed primers. (A) Primer Pair C at 60–65°C, (B) Primer Pairs D and E at 48–62°C.

Primer C produced 7 amplicons, whereas Primer D and Primer E produced 10 and 11 amplicons, respectively (Table 3). Primer E amplified the same samples as Primer D with the addition of Sample 4 and 12, but it was unable to amplify Sample 5. Primer E amplified the same samples as Primer C except for Samples 5 and 19, which were amplified only by Primer C. Only Primer E was able to amplify Sample 6. After DNA sequencing, however, Sample 6 exhibited unreadable sequence results. A BLAST analysis of the 13 readable DNA sequences verified that the they were specific to *Fragaria ANR*. The 134-bp long DNA sequences were then used for SNP discovery.

**Table 3.** SNP profiles of the 13 cultivar specimens based on the full array of 11 SNP markers.

| Cultivars | SNP ID |  |  |  |  |  |  |  |  |  |  |
| --- | --- | --- | --- | --- | --- | --- | --- | --- | --- | --- | --- |
|  | 1 | 2 | 3 | 4 | 5 | 6 | 7 | 8 | 9 | 10 | 11 |
| Sweet Charlie Arg2 | -- | TT | GG | AA | -- | AA | CC | CC | AA | AA | AA |
| Sweet Charlie Arg3 | -- | TT | GG | AA | -- | AA | CC | CC | AA | AA | AA |
| Sweet Charlie Wash1 | -- | TT | GG | AA | -- | AA | CC | CC | AA | AA | AA |
| Sweet Charlie4 | -- | TT | GG | AA | -- | AA | CC | CC | AA | AA | AA |
| Strawberry Festival1 | -- | TT | GG | AA | -T | AA | CC | CC | AA | AA | AA |
| Sweet Charlie2 | -- | TT | GG | AA | -T | AA | CC | CC | AA | AA | AA |
| Strawberry Festival3 | -- | TC | G- | AG | -A | AC | CT | CA | AA | AA | AA |
| Sweet Charlie Wash2* | -- | TC | GT | AG | -A | AC | CT | CA | AA | AA | AA |
| Sweet Charlie6 | -- | TC | GT | AG | -A | AC | CC | CC | AA | AA | AA |
| Winterdawn2 | -- | TC | GT | AG | -A | AC | CC | CC | AA | AA | AA |
| Winterdawn3 | -- | TC | GT | AG | -A | AC | CC | CC | AA | AA | AA |
| Sweet Charlie California | -- | TT | GG | AA | -- | AA | CC | CC | AA | AA | AC |
| Strawberry Festival2 | -- | TC | GG | AA | -C | AA | CT | CC | AA | AG | AC |

\*Wash = Washington

The use of real-time PCR followed by MCA with fluorescent dyes has proven to be a highly sensitive method in the discovery of mutations and the genotyping of SNPs [27]. This combination is a fast, simple, and low-cost method for SNP genotyping that is nonetheless precise, reproducible, and flexible with a high-throughput capability. Its uses have been described for humans, animals, and plants [29,30,33–37]. Although high-resolution MCA is a more powerful tool for SNP characterization, standard MCA has been shown to be sufficient for SNP genotyping wherein each melting curve correlates with the specificity of genotype reactions [36,37]. In this study, MCA was performed to ensure reaction specificity of the primers for SNP detection. The MCA results showed successful amplification of 14 out of 20 strawberry samples. The successful amplicons were further verified by visualizing using agarose gel electrophoresis. Although gel electrophoresis is an unnecessary step after real-time PCR and MCA, discrepancies between melt peak data and gel electrophoresis bands have been reported in other crops [38,39]. But such discrepancies were not detected in this study, with melt peak data coinciding with amplicons visualized by gel electrophoresis (data not shown).

### 3.3. SNP detection and cluster analysis

Alignment of the 13 DNA sequences produced 11 SNP markers (Table 4). Eight of the SNPs were biallelic and three were triallelic. There were three instances of transversions, three transitions, and four InDels. The 11 SNP markers were then used for cluster analysis. The markers were able to discriminate the samples into five distinct genotypic clusters with 7 distinctive genotypes (Table 4). ‘Sweet Charlie’ 4, ‘Sweet Charlie’ Argentina 2, ‘Sweet Charlie’ Argentina 3, and ‘Sweet Charlie’ Washington 1 had the same SNP profiles. ‘Sweet Charlie’ 2 and ‘Strawberry Festival’ 1 had the same SNP profile as the previous four except for one insertion at SNP5. In another cluster, ‘Strawberry Festival’ 3 differed from the similarly clustered ‘Sweet Charlie’ Washington 2 due to an insertion at SNP3. ‘Sweet Charlie’ 6, ‘Winter Dawn’ 2, and ‘Winter Dawn’ 3 had the same SNP profiles. ‘Sweet Charlie’ California differed in its SNP profile from the ‘Sweet Charlie’ samples of the first cluster due to an insertion at SNP11. Finally, in the last cluster, ‘Strawberry Festival’ 2 differed from the previous two due to an insertion at SNP5.

The use of SNP markers for genotypic characterization has become widespread in recent years for many diploid crops. In the strawberry, the discovery of SNPs has not been as easily accomplished due to the complexity posed by its allo-octoploid nature. The challenges posed by polyploid crops to SNP discovery have been highlighted in several published papers, most notably by Bundock et al, Kaur et al., Clevenger et al., Garcia et al., and Mammadov et al. [21–23,40,41]. Kaur et al. and Clevenger et al. suggested some strategies for overcoming the difficulties in developing SNP markers for polyploids [22,23]. Following such strategies, this study used sequence information from EST libraries, focused SNP discovery on a single gene, used the CAP3 DNA Sequence Assembler for primer discovery, and employed MCA and Sanger sequencing for SNP validation.

Many molecular markers for the cultivated strawberry have been developed for their potential use in cultivar identification and genetic diversity studies [4]. For instance, Kunihisa et al. successfully verified Japanese cultivars obtained from a supermarket using CAPS markers [9]. In the United States, SSR markers, such as those developed by Dangl et al. and Lewers et al. [4,42], are currently the markers of choice in both public and private laboratories. Majority of the samples sent for verification are from nurseries, with the rest coming from breeding programs [4]. The main rationale for cultivar identification is to prevent misidentification of planting materials in nurseries, especially after several cycles of runner propagation, protecting the intellectual property rights of plant breeders, and ensuring correct identification of germplasm collections [4,9]. Molecular markers have already been used to settle cases of patent infringement in certain strawberry cultivars [12,43].

The utilization of SNP markers in the strawberry is quite new. Ge et al. (2013) were first to report on the use of SNP markers to discriminate among cultivars in China [44]. But the problems associated with strawberry polyploidy were not tackled in their report. The IStraw90, a 90 K Axiom^®^ SNP array is currently the most comprehensive system for SNP detection and verification for this crop [45]. Sargent et al. used the IStraw90 to create a linkage map containing SNP markers [46], whereas Nagano et al. used a combination of candidate SNPs and SSRs to create a linkage map using the IStraw90 [47]. However, use of the array is expensive at US$80 to US$105 per sample [48]. Verma et al. developed a less expensive protocol based on the IStraw90 at US$50 per sample [48]. Jung et al. also developed another SNP genotyping system, the Fluidigm 24 SNPs genotyping system, based on the IStraw90 [49]. The cost for SNP detection and marker development using these systems is prohibitive for strawberry-producing developing countries such as the Philippines that want to utilize SNPs for MAS and cultivar verification. In this study, we were able to develop a simple, fast, and inexpensive protocol for SNP discovery and utilization in cultivar verification. Although this method does not provide a wide-ranging and thorough study of SNPs in the strawberry, it has practical applications and requires only standard molecular biology equipment and freely available software.

### 3.4. Cluster analysis

A phylogenetic tree was subsequently constructed using the Neighbor-Joining method based on the SNP marker information (Fig. 3). Cluster analysis grouped the strawberry cultivars into five clusters. Cluster 1, the largest, included five ‘Sweet Charlie’ samples and one ‘Strawberry Festival’ sample. Cluster 2 had only one ‘Sweet Charlie’ sample, this time from California. Cluster 3 had one ‘Strawberry Festival’ sample. Cluster 4 included the ‘Sweet Charlie’ sample from Washington (state, USA) and one ‘Strawberry Festival’ sample. Lastly, Cluster 5 included one ‘Sweet Charlie’ sample and the two ‘Winter Dawn’ samples.

**Figure 3.**
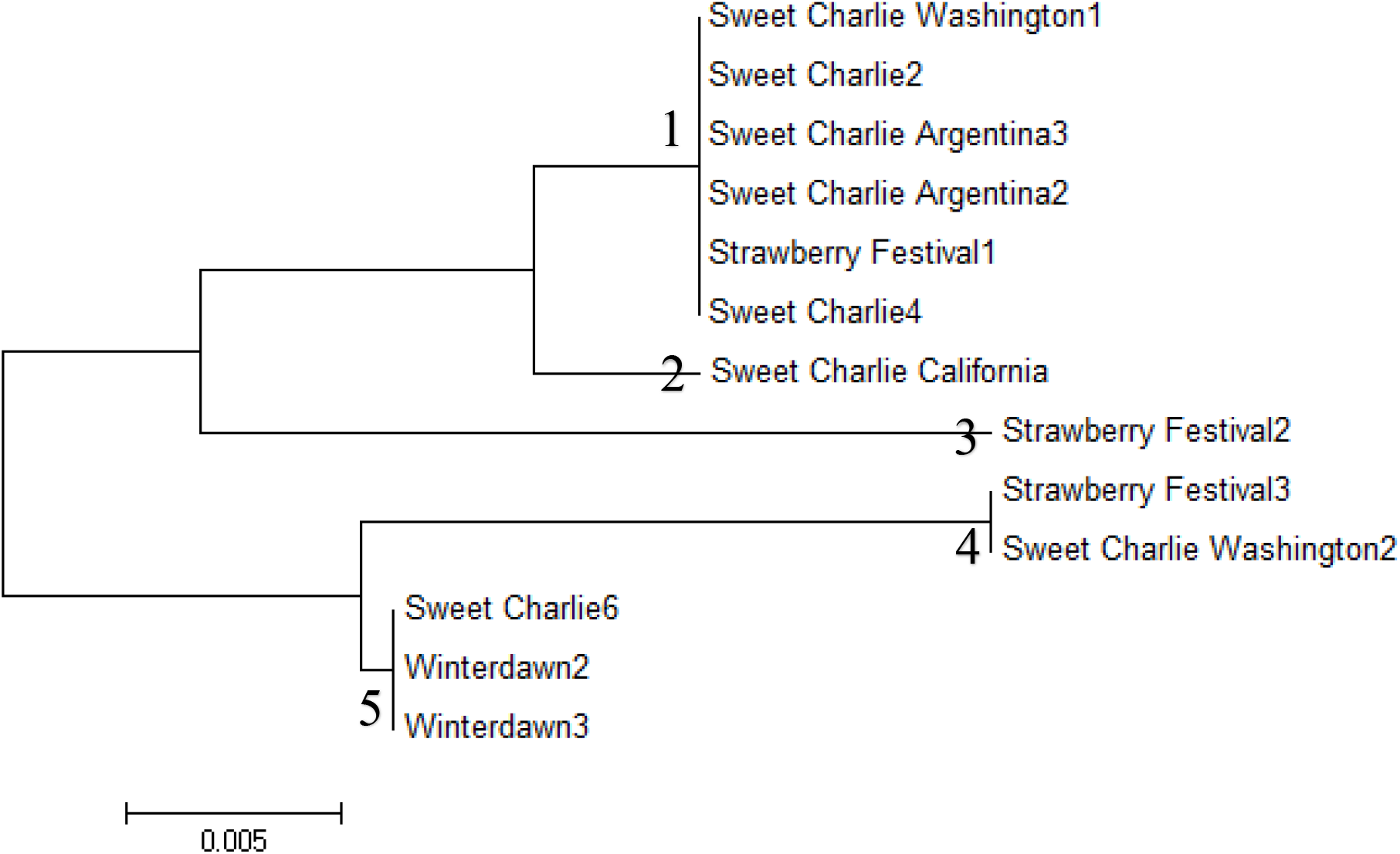
Phylogenetic tree of 13 strawberry cultivars collected at the Strawberry Farm

One easily noticeable revelation from the phylogenetic tree is the clustering of the different ‘Sweet Charlie’ specimens into separate genotypic groups. Five of the ‘Sweet Charlie’ specimens were grouped in Cluster 1 and one specimen each were placed in Clusters 2, 4, and 5. It appears that some of these supposedly ‘Sweet Charlie’ specimens were not properly identified by the farmers. The same could be said for the three ‘Strawberry Festival’ specimens, which were clustered into separate genotypic groups (Clusters 1, 3, and 4). On the other hand, the two ‘Winter Dawn’ specimens were grouped in the same cluster (Cluster 5), indicating correct identification for both.

The exact location at the Farm where each of these cultivars were collected is shown in Fig. 4. In Sampling Area 3, the farmers claimed to grow two cultivars only: ‘Sweet Charlie’ and ‘Strawberry Festival’. However, the three ‘Sweet Charlie’ samples obtained from this location, which had different origins, belonged to different clusters. Furthermore, the cultivar that they identified as ‘Strawberry Festival’ is clustered with ‘Sweet Charlie’ Washington2. The same could be said for Sampling Area 1 where two ‘Strawberry Festival’ samples growing in the same plot had different SNP profiles (Table 3). The constant exchange of planting materials among farmers may have already resulted in a new mix-up wherein the ‘Winter Dawn’ plants in Sampling Area 6 and ‘Sweet Charlie’6 in Sampling Area 2 had identical SNP profiles (Table 4 and Fig. 4).

**Figure 4.**
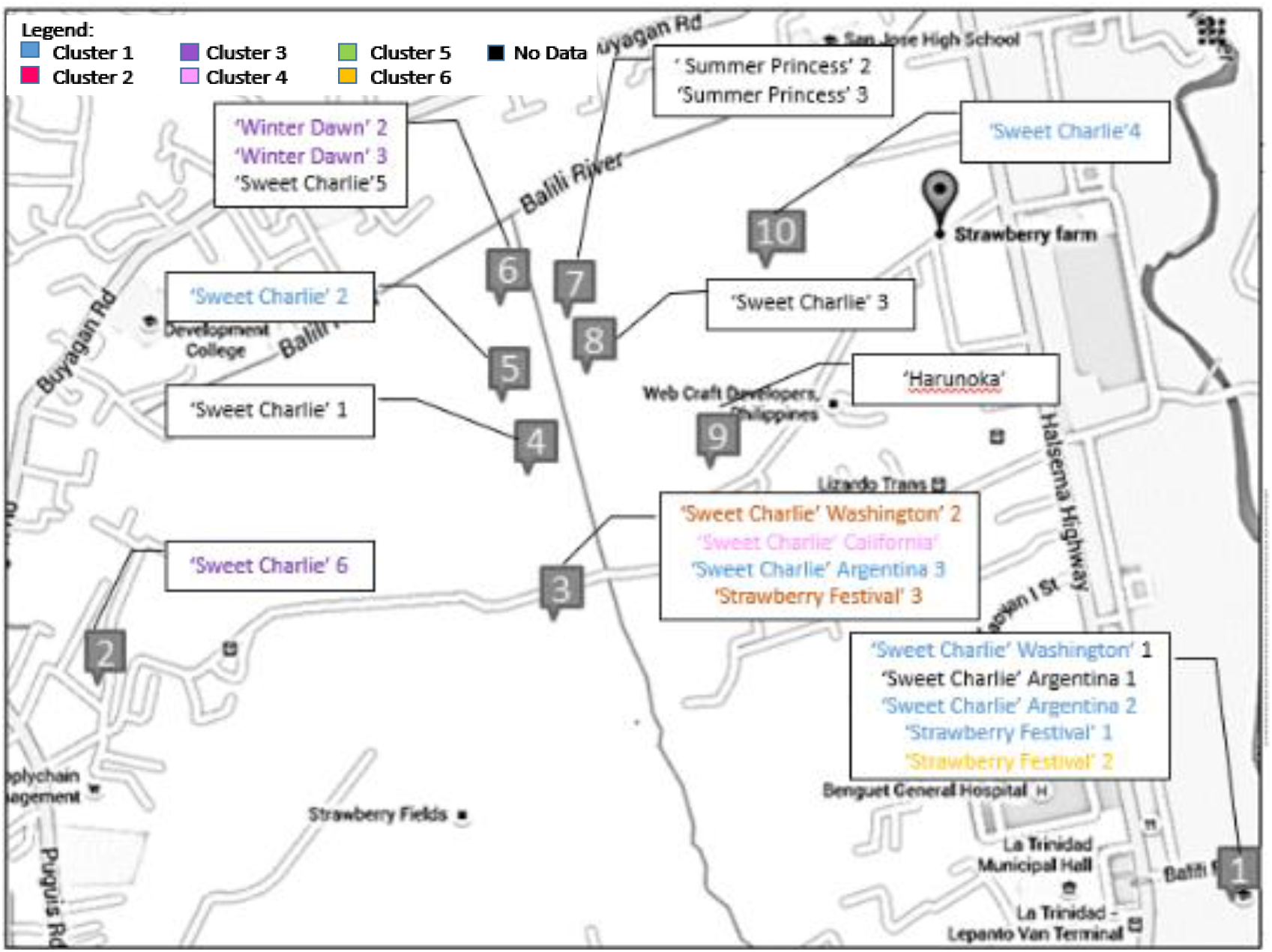
Map showing where and which strawberry cultivars were collected per sampling area. Colors indicate the Cluster to which the cultivars belong based on SNP profiles.

Strawberry cultivars in the Strawberry Farm are mainly propagated by cloning, either through plant tissue culture or the rooting of runners, so different plants identified as a certain cultivar should have the same genotypic identities. But the clustering analysis did not bear out the farmers’ identification of the cultivars they were planting. This raises the concern for the correct identity of the strawberries being produced in the area.

There are very few reports on misidentification of strawberry cultivars based on molecular marker information. Garcia et al. discovered three accessions of ‘Pajaro’, obtained from the field, that exhibited different amplification profiles based on RAPD markers [7]. In this study, the results indicated misidentification in the field where plants grown on the same plot and identified as one cultivar turned out to be genetically distinct from each other. Conversely, samples that were genotypically identical were identified as distinct cultivars (Figs. 3 and 4). Cases of cultivar misidentification as detected by molecular markers have also been reported in blueberry [50], apple [51], and black walnut [52].

The large grouping of plant samples in Cluster 1 on the phylogenetic tree (Fig. 3) could be due to several reasons. It is highly likely that some of these samples are misidentified. There is also a possibility that cultivars may have been misidentified at the country of origin. Another reason could be that these cultivars share very close genetic relationships based on their sequence alignments (Table 4). It is also possible that the large grouping indicates that the SNP markers used, which were developed from a single gene, were not enough to discriminate the samples from each other. The effect of choosing molecular markers based on a specific breeding trait can also influence clustering. Gil-Ariza et al. reported that tested cultivars grouped according to breeding period and specific climatic adaptation traits [53]. It is not clear which among the ‘Sweet Charlie’ samples analyzed in the study is the true ‘Sweet Charlie’. But the identical sequences of the ‘Sweet Charlie’ samples in Cluster 1 indicate that they came from the same genetic source. Difficulties in distinguishing cultivars at the subspecies level is challenging for single genes, an observation highlighted in the case of grapevine [54]. But the complexity of the strawberry genome also means that in order to develop SNP markers for a specific purpose (i.e., verifying misidentifications), using a single gene may be the more appropriate choice [22].

### 3.5. Leaf morphology

Figure 5 shows that although all of the leaf samples collected were supposedly of ‘Sweet Charlie,’ the molecular analysis revealed that Samples A, B, D, F, and G belong to Cluster 1. On the other hand, Samples C, E, and H belong to Clusters 5, 4, and 2, respectively (Table 4). Morphologically speaking, the leaf shape (terminal and side leaflets) of Sample B is different from the other samples classified in Cluster 1. In fact, it is more similar to Sample C. The leaf of ‘Sweet Charlie’ is described as obovate, with the terminal leaflet having an obtuse base and the side leaflets having oblique bases with rounded dentate margins [55]. Based on this description, for samples in Cluster 1, Samples D and F have obtuse terminal leaflet bases, whereas Samples A, B, and G have round bases. All of the leaf samples have side leaflets with oblique bases. Only Samples D, F, and G can be described as having round-dentate leaf margins, whereas Samples A and B have dentate margins.

Of all the leaf samples shown in Fig. 5, Samples D and F closely match the leaf descriptions of ‘Sweet Charlie’ in Cluster 1. However, Sample E (Cluster 4) and Sample H (Cluster 2) also match this description. Although strawberry leaves can vary in other aspects such as gloss and color of the adaxial and abaxial sides [56], it is difficult to assess the leaves based on such characteristics because they are traits most prone to environmental influence such as the availability of nutrients. Because the leaf samples were taken from different farmers, cultural management practices, especially fertilization, can greatly differ, leading to differences in leaf color and glossiness even in the same cultivar.

**Figure 5.**
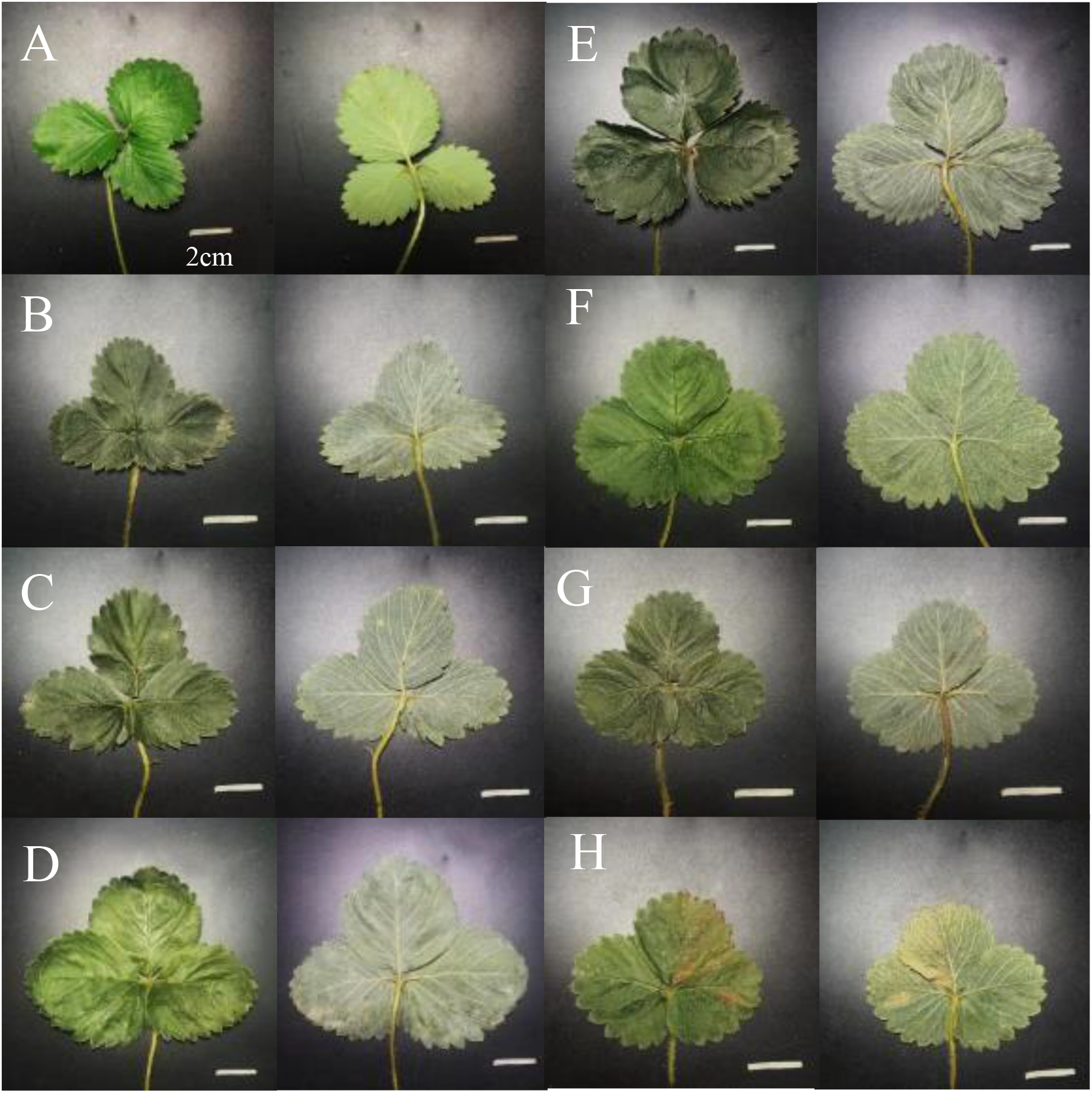
Comparison of adaxial (left) and abaxial (right) leaf surfaces of the different ‘Sweet Charlie’ samples collected at the Strawberry Farm. (A) ‘Sweet Charlie’ 2, (B) ‘Sweet Charlie’ 4, (C) ‘Sweet Charlie’ 6, (D) ‘Sweet Charlie’Washington 1, (E) ‘Sweet Charlie’ Washington 2, (F) ‘Sweet Charlie’Argentina 2, (G) ‘Sweet Charlie’ Argentina 3, and (H) ‘Sweet Charlie’ California.

The leaf samples illustrate the great difficulty in characterizing strawberry plants especially at the juvenile stage when they are mass propagated and transported to farmers. Characterizing strawberry cultivars using morphological features are, therefore, not the most reliable method to distinguish them from each other. This is true even for other crops because physiological and environmental influences can modify these traits [5].

Without the benefit of molecular tools, farmers rely solely on morphological characteristics, particularly the leaves, when differentiating cultivars. But leaves of young strawberry plants of different cultivars can appear nearly identical. And even leaves from the same cultivar can appear different from each other. If there are fine distinctions, these are often missed by the farmers who are not trained on morphological characterizations.

The case of misidentification for ‘Sweet Charlie’ is of particular importance to the Philippine strawberry industry. A sequence of ‘Sweet Charlie’ was included in the BLAST search to ensure that an EST of the cultivar was included due to it being the most propagated in the Philippines. Thus, verification of the proper identity of this cultivar is of high importance. All accessions of ‘Strawberry Festival’ and ‘Sweet Charlie’ were collected from the Strawberry Farm, and there are no certified specimens for these two cultivars in the country that can be used for molecular verification. Thus, it could not be determined which of the strawberry samples is the genuine cultivar. The results of this study could only elucidate the relationships between the samples collected based on similar or different genotypic identities. Further studies are thus needed to clarify this point, not just for ‘Sweet Charlie’ but also for the other misidentified cultivars. A larger study also needs to be conducted to determine the extent of misidentifications among other cultivars being grown in the Philippines.

In conclusion, we developed 11 SNP markers from the *ANR* gene of the cultivated strawberry to detect misidentifications among cultivars grown in La Trinidad, Benguet, Philippines. The methods used in this verification study is of practical importance to the strawberry industry because it is a relatively inexpensive, simple, fast, sensitive, and high-throughput, single-tube assay for SNP detection. It requires only a DNA extraction kit, a PCR Kit, a real-time PCR system, freely available software, and the cost of DNA sequencing. The results of this study indicate that the farmers’ identification of strawberry cultivars was not in line with the molecular evidence, which revealed that many of the plant samples tested were inaccurately identified based on clustering analysis.

## Acknowledgement

The authors would like to sincerely thank Mr. Jordan Tomin and Dr. Danilo Padua of Benguet State University for assisting in the strawberry collection and the laboratory staff of UP Manila National Institutes of Health for allowing the authors to use their lab for the molecular portion of the study.

## References

[1] Food and Agriculture Organization of the United Nations [homepage on the Internet]. Statistical databases: Crops. 2019 [cited 2019 April 5]. Available from: http://www.fao.org/faostat/en/#data/QC (2019).

[2] Darrow GM. The strawberry: history, breeding and physiology. New York: Holt, Rinehart and Winston; 1966.

[3] Fernandez RA. Strawberry industry headed for better times – study. Philippine Star [homepage on the Internet]. 2016 [cited Available from: http://www.philstar.com/business-usual/713975/strawberry-industry-headed-better-times-study (2016).

[4] Whitaker VM. Applications of molecular markers in strawberry. J Berry Res. 2011; 1: 115–127. doi:10.3233/BR-2011-013.

[5] Dangl GS, Lee EW, Sim ST, Golino DA. A new system for strawberry cultivar identification developed at Foundation Plant Services (FPS), University of California, Davis, using simple sequence repeat (SR) primers. NASS/NASGA Proc. 2007; 118–121.

[6] Bell JA, Simpson DW. The use of isoenzyme polymorphisms as an aid for cultivar identification in strawberry. Euphytica 1994; 77:113–17.

[7] Garcia MG, Ontivero M, Diaz Ricci JC, Castagnaro A. Morphological traits and high resolution RAPD markers for the identification of the main strawberry varieties cultivated in Argentina. Plant Breed. 2002; 121:76–80.

[8] Brunings AM, Moyer C, Peres N, Folta KM. Implementation of simple sequence repeat markers to genotype Florida strawberry varieties. Euphytica 2010; 173:63–75. doi:10.1007/s10681-009-0112-4.

[9] Kunihisa M, Fukino N, Matsumoto S. Development of cleavage amplified polymorphic sequence (CAPS) markers for identification of strawberry cultivars. Euphytica 2003; 134:209–15.

[10] Gidoni D, Rom M, Kunik T, Zur M, Izsak E, Izhar S, et al. Strawberry-cultivar identification using randomly amplified polymorphic DNA (RAPD) markers. Plant Breed. 1994; 113:339–42.

[11] Degani C, Rowland LJ, Levi A, Hortynski JA, Galletta GJ. DNA fingerprinting of strawberry *(Fragaria* x *ananassa)* cultivars using randomly amplified polymorphic DNA (RAPD) markers. Euphytica 1998; 102:247–53.

[12] Congiu L, Chicca M, Cella R, Rossi R, Bernacchia G. The use of random amplified polymorphic DNA (RAPD) markers to identify strawberry varieties: a forensic application. Mol Ecol. 2000; 9:229–32.

[13] Degani C, Rowland LJ, Saunders JA, Hokanson SC, Ogden EL, et al. A comparison of genetic relationship measures in strawberry *(Fragaria* × *ananassa* Duch.) based on AFLPs, RAPDs, and pedigree data. Euphytica 2001; 117:1–12.

[14] Tyrka M, Dziadczyk P, Hortyinski JA. Simplified AFLP procedure as a tool for identification of strawberry cultivars and advanced breeding lines. Euphytica 2002; 125:273–80.

[15] Peng M, Zong X, Wang C, Meng F. Genetic diversity of strawberry *(Fragaria ananassa* Duch.) from the Motuo County of the Tibet Plateau determined by AFLP markers. Biotech Biotech Equip. 2015; 29:876–81. doi:10.1080/13102818.2015.1050968.

[16] Honjo M, Nunome T, Kataoka S, Yano T, Yamazaki H, Hamano M, et al. Strawberry cultivar identification based on hypervariable SSR markers. Breed Sci. 2011; 61:420–25. doi:10.1270/jsbbs.61.420.

[17] Lim S, Lee J, Lee HJ, Park KH, Kim DS, Min SR, et al. The genetic diversity among strawberry breeding resources based on SSRs. Sci Agric. 2017; 74:226–34. doi:10.1590/1678-992X-2016-0046.

[18] Rugienius R, Šikšnianienė JB, Frercks B, Stanienė G, Stepulaitienė I, Haimi P, et al. Characterization of strawberry *(Fragaria* × *ananassa* Duch.) cultivars and hybrid clones using SSR and AFLP markers. Zemdirbyste-Agriculture 2015; 102:177–84. doi:10.13080/z-a.2015.102.023.

[19] Günaydın S, Kafkas S. Characterization of strawberry cultivars by SSR and CAPS markers. Acta Hortic. 2017; 1156:171–77. doi:10.17660/ActaHortic.2017.1156.25.

[20] Wada T, Noguchi Y, Isobe S, Kunihisa M, Sueyoshi T, Shimomura K, et al. Development of a core collection of strawberry cultivars based on SSR and CAPS marker polymorphisms. Hortic J. 2017; 86:365–78. doi:10.2503/hortj.MI-142.

[21] Bundock PC, Eliott, FG, Ablett G, Benson AD, Casu RE, Aitken KS, et al. Targeted single nucleotide polymorphism (SNP) discovery in a highly polyploid plant species using 454 sequencing. Plant Biotech J. 2009; 7:347–54. doi:10.1111/j.1467-7652.2009.00401.x.

[22] Kaur S, Francki MG, Forster JW. Identification, characterization and interpretation of single-nucleotide sequence variation in allopolyploid crop species. Plant Biotech J. 2012; 10:125–38. doi:10.1111/j.1467-7652.2011.00644.x.

[23] Clevenger J, Chavarro C, Pearl SA, Ozias-Akins P, Jackson SA. Single nucleotide polymorphism identification in polyploids: a review, example, and recommendations. Mol Plant. 2015; 8:831–46. doi:10.1016/j.molp.2015.02.002.

[24] Hawkins C, Caruana J, Schiksnis E, Liu Z. Genome-scale DNA variant analysis and functional validation of a SNP underlying yellow fruit color in wild strawberry. Sci Rep. 2016; 6:29017. doi:10.1038/srep29017.

[25] Mahoney LL, Sargent DJ, Abebe-Akele F, Wood DJ, Ward JA, Bassil NV, et al. A high-density linkage map of the ancestral diploid strawberry, *Fragaria iinumae*, constructed with single nucleotide polymorphism markers from the IStraw90 Array and genotyping by sequencing. Plant Genom. 9. doi:10.3835/plantgenome2015.08.0071.

[26] Han Y, Kang Y, Torres-Jerez I, Cheung F, Town CD, Zhao PX, et al. Genome-wide SNP discovery in tetraploid alfalfa using 454 sequencing and high resolution melting analysis. BMC Genom. 2011; 12:350. doi:10.1186/1471-2164-12-350.

[27] Han Y, Khu DM, Monteros MJ. High-resolution melting analysis for SNP genotyping and mapping in tetraploid alfalfa *(Medicago sativa* L.). Mol Breed. 2012; 29:489–501. doi:10.1007/s11032-011-9566-x.

[28] Li X, Acharya A, Farmer AD. Prevalence of single nucleotide polymorphism among 27 diverse alfalfa genotypes as assessed by transcriptome sequencing. BMC Genom. 2012; 13:568. doi:10.1186/1471-2164-13-568.

[29] Baris I, Etlik O, Koksal V, Ocak Z, Baris ST. SYBR green dye-based probe-free SNP genotyping: Introduction of T-Plex real-time PCR assay. Anal Biochem. 2013; 441:225–31. doi:10.1016/j.ab.2013.07.007.

[30] Pryor RJ, Wittwer CT. Real-time polymerase chain reaction and melting curve analysis in Methods in molecular biology 336: clinical applications of PCR. (eds Lo, Y. M. D., Chiu, R. W. K. & Chan, K. C. A.) 19–32 (Humana Press Inc., NJ, 2006).

[31] Almeida JRM, D’Amico E, Preuss A, Carbone F, de Vos RCH, Deiml B, et al. Characterization of major enzymes and genes involved in flavonoid and proanthocyanidin biosynthesis during fruit development in strawberry *(Fragaria* × *ananassa)*. Arch Biochem Biophys. 2007; 465:67–71. doi:10.1016/j.abb.2007.04.040.

[32] Carbone F, Preuss A, De Vos RCH, D’amico E, Perrotta G, Bovy AG, et al. Developmental, genetic and environmental factors affect expression of flavonoid genes, enzymes and metabolites in strawberry fruits. Plant, Cell Environ. 2009; 32:1117–31. doi:10.1111/j.1365-3040.2009.01994.x.

[33] Lipsky RH, Mazzanti CM, Rudolph JG, Xu K, Vyas G, Bozak D, et al. DNA melting analysis for detection of single nucleotide polymorphisms. Clin Chem. 2001; 47:635–44.

[34] Papp AC, Pinsonneault JK, Cooke G, Sadée W. Single nucleotide polymorphism genotyping using allele-specific PCR and fluorescence melting curves. BioTechniques. 2003;34:1068–72.

[35] Liew M, Pryor R, Palais R, Meadows C, Erali M, Lyon E, et al. Genotyping of single-nucleotide polymorphisms by high-resolution melting of small amplicons. Clin Chem. 2004; 50:1156–64.

[36] Zhou L, Myers AN, Vandersteen JG, Wang L, Wittwer CT. Closed-tube genotyping with unlabeled oligonucleotide probes and a saturating DNA dye. Clin Chem. 2004; 50:1328–35.

[37] Varga A, James D. Real-time RT-PCR and SYBR Green I melting curve analysis for the identification of plum pox virus strains C, EA, and W: Effect of amplicon size, melt rate, and dye translocation. J Virol Method. 2005; 132:146–53. doi: 10.1016/j.jviromet.2005.10.004.

[38] Giglio S, Monis PT, Saint CP. Demonstration of preferential binding of SYBR Green I to specific DNA fragments in real-time multiplex PCR. Nucleic Acids Res. 2003; 31:e136. doi:10.1093/nar/gng135.

[39] Monis PT, Giglio S, Saint CP. Comparison of SYTO9 and SYBR Green I for real-time polymerase chain reaction and investigation of the effect of dye concentration on amplification and DNA melting curve analysis. Anal Biochem. 2005; 340:24–34.

[40] Garcia AAF, Mollinari M, Marconi TG, Serang OR, Silva RR, Vieira MLC, et al. SNP genotyping allows an in-depth characterisation of the genome of sugarcane and other complex autopolyploids. Sci Rep. 2013; 3:3399. doi:10.1038/srep03399.

[41] Mammadov J, Aggarwal R, Buyyarapu I, Kumpatla S. SNP markers and their impact on plant breeding. Int J Plant Genom. 2012; 12. doi:10.1155/2012/728398.

[42] Lewers KS, Styan SMN, Hokanson SC. Strawberry GenBank-derived and genomic simple sequence repeat (SSR) markers and their utility with strawberry, blackberry, and red and black raspberry. J Am Soc Hortic Sci. 2005; 130:102–115.

[43] Chivvis M. Strawberry fields. Intellectual Property Magazine. 2017. November.

[44] Ge AJ, Han J, Li XD, Zhao MZ, Liu H, Dong QH, et al. Characterization of SNPs in strawberry cultivars in China. Gen Mol Res. 2013; 12:639–45.

[45] Bassil NV, Davis TM, Zhang H, Ficklin S, Mittmann M, Webster T, et al. Development and preliminary evaluation of a 90 K Axiom^®^ SNP array for the allo-octoploid cultivated strawberry *Fragaria* × *ananassa*. BMC Genom. 2015; 16:15. doi:10.1186/s12864-015-1310-1.

[46] Sargent DJ, Yang Y, Surbanovski N, Bianco L, Buti M, Velasco R, et al. HaploSNP affinities and linkage map positions illuminate sub-genome composition in the octoploid, cultivated strawberry *(Fragaria* × *ananassa)*. Plant Sci. 2016; 242:140–50. doi:10.1016/j.plantsci.2015.07.004.

[47] Nagano S, Shirasawa K, Hirakawa H, Maeda F, Ishikawa M, Isobe SN. Discrimination of candidate subgenome-specific loci by linkage map construction with an S1 population of octoploid strawberry *(Fragaria* × *ananassa)*. BMC Genom 2017; 18:374. doi:10.1186/s12864-017-3762-y.

[48] Verma S, Bassil NV, van de Weg E, Harrison RJ, Monfort A, Hidalgo JM, et al. Development and evaluation of the Axiom^®^ IStraw35 384HT array for the allo-octoploid cultivated strawberry *Fragaria* × *ananassa*. Acta Hortic. 2017; 1156:75–81. doi:10.17660/ActaHortic.2017.1156.10.

[49] Jung HJ, Veerappan K, Natarajan S, Jeong N, Hwang I, Nagano S, et al. A system for distinguishing octoploid strawberry cultivars using high-throughput SNP genotyping. Tropical Plant Biol. 2017; 10:68–76. doi:10.1007/s12042-017-9185-8.

[50] Hinrichsen P, Herminia Castro M, Ravest G, Rojas G, Mendez M, Bassil NV, et al. Minimal microsatellite marker panel for fingerprinting blueberry cultivars. Acta Hortic. 2009; 810:173–80.

[51] Storti A, Via JD, Baric S. Comparative molecular genetic analysis of apple genotypes maintained in germplasm collections. Erwerbs-Obstbau 2012; 54:137–41. doi:10.1007/s10341-012-0168-5.

[52] Zhao P, Zhou H, Coggeshall M, Reid B, Woeste K. Discrimination and assessment of black walnut *(Juglans nigra* L.) cultivars using phenology and microsatellite markers (SSRs). Can J Plant Sci. 2018; 98:616–27. doi:10.1139/cjps-2017-0214.

[53] Gil-Ariza DJ, Amaya I, Lopez-Aranda JM, Sanchez-Sevilla JF. Impact of plant breeding on the genetic diversity of cultivated strawberry as revealed by expressed sequence tag-derived simple sequence repeat markers. J Am Soc Hortic Sci. 2009; 134:337–47.

[54] Nicolè S, Barcaccia G, Erickson DL, Kress JW, Lucchin M. The coding region of the UFGT gene is a source of diagnostic SNP markers that allow single-locus DNA genotyping for the assessment of cultivar identity and ancestry in grapevine *(Vitis vinifera* L.). BMC Res Notes. 2013; 6:502. doi:10.1186/1756-0500-6-502.

[55] Howard CM. United States Patent No. USPP8729P (1994).

[56] Höfer M, Drewes-Alwarez R, Scheewe P, Olbricht K. Morphological evaluation of 108 strawberry cultivars—and consequences for the use of descriptors. J Berry Res. 2012; 2:191–206. doi:10.3233/JBR-2012-042.

